# Thermodynamic analysis of mitochondrial DNA breakpoints reveals mechanistic details of deletion mutagenesis

**DOI:** 10.1101/254631

**Authors:** Lakshmi Narayanan Lakshmanan, Zhuangli Yee, Jan Gruber, Barry Halliwell, Rudiyanto Gunawan

**Affiliations:** Institute for Chemical and Bioengineering, ETH Zurich, Zurich 8093, Switzerland; Swiss Institute of Bioinformatics, Quartier Sorge – Batiment Genopode, 1015 Lausanne, Switzerland; Department of Biochemistry, Yong Loo Lin School of Medicine, National University of Singapore, Singapore; Ageing Research Laboratory, Science Division, Yale-NUS College, Singapore

**Author notes:** Corresponding Author, Address: Rudiyanto Gunawan, Institute for Chemical and Bioengineering, ETH Zurich, Vladimir-Prelog-Weg 1, Zurich 8093, Switzerland.

**Keywords:** Mitochondrial DNA, deletion mutation, double-strand breaks, DNA hybridization, thermodynamics, DSB repair, strand invasion

## Abstract

Broad evidence support double-strand breaks (DSBs) as initiators of mitochondrial DNA (mtDNA) deletion mutations. But the mechanism of DSB-induced deletions, including the DSB repair pathway(s) involved, remains to be established. Here, we used DNA hybridization thermodynamics to analyze misalignment lengths surrounding deletion breakpoints. Our analysis of 9,655 previously reported mammalian mtDNA deletions and 1,307 novel *Caenorhabditis elegans* mtDNA deletions, indicates a significant role of 0–25bp misalignments, supporting the role of erroneous non-homologous and micro-homology dependent DSB repair in deletion formation. Based on these insights we propose that DSB-induced mtDNA deletions occur via the misjoining of DSB ends and/or strand invasion of open mtDNA regions by DSB ends.

---

Pathogenic deletion mutations of mitochondrial DNA (mtDNA) typically involve the loss of several hundreds to thousands of nucleotides encoding key proteins involved in the mitochondrial electron transport chain and ATP synthesis. The accumulation of such mutant mtDNA molecules in a cell leads to mitochondrial dysfunction and cellular energy crisis. In humans, mtDNA deletion mutations cause mitochondrial diseases such as Kearns-Sayre Syndrome (KSS)^1,2^, Chronic Progressive External Ophthalmoplegia (CPEO)^1,2^ and Pearson Syndrome (PS)^3,4^, and have been implicated in age-related diseases such as sarcopenia^5^. Mitochondrial DNA deletions have also been reported in tissues of patients suffering from Parkinson’s disease^6^, epilepsy^7^, myositis^8^, Charcot Marie Tooth Disease (CMTD)^9^, diabetes^10^, and cancer^11^ although their role in these diseases is less clearly defined.

The mechanism of mtDNA deletion formation is of fundamental importance for understanding loss of mtDNA integrity and has been a subject of great interest and debate^12^. A commonly reported feature of mtDNA deletions is the presence of direct repeat (DR) sequences flanking the deletion breakpoints^13–16^. This observation has led to two major models of mtDNA deletion formation, namely the “slip-strand” model and the “double-strand break (DSB) repair error” model. The slip-strand model hypothesizes that mtDNA deletions occur during mtDNA replications, more specifically involving the strand-displacement mode of replication^17^. In strand-displacement replication, the syntheses of the heavy and light mtDNA strands are not synchronized with the parental heavy strand being displaced during the replication of the leading strand. The displaced strand remains single-stranded until roughly 65% of the daughter heavy strand is synthesized. Such single-stranded heavy strand is prone to misalignments (i.e. hybridization of DNA sequences from different positions) with the exposed light strand, an error that involves DR sequences, leading to the formation of a deletion mutation^17^. But, this slip-strand hypothesis has recently been put into question^12^. One of the main objections is that the slip-strand model requires the presence of “naked” single-stranded mtDNA regions during replication for DNA misalignments to occur. However, studies on mtDNA replication have demonstrated that the lagging-strand of mtDNA is in fact protected by mitochondrial single strand binding proteins or bound by RNA molecules and is therefore not as free and open as previously thought^18^.

Meanwhile, the DSB repair error model, like the name suggests, attributes mtDNA deletions to erroneous repair of DSBs. Direct support for DSBs as an initiating step in mtDNA deletion formation comes from transgenic mouse models expressing mitochondria-targeted restriction endonuclease^19–21^. Mitochondrial expression of these mtDNA targeting enzymes causes DSBs reliably at specific sequence locations (restriction sites) in the mtDNA. In these mouse models, induction of DSBs by this method promotes the formation of mtDNA deletions with breakpoints near the restriction sites. Endogenous and exogenous DNA insults that induce or promote DSBs, such as oxidative stress and ionizing radiation have also been shown to increase the occurrence of mtDNA deletions^7,22^. Furthermore, there exist broad experimental evidence that mtDNA replication fork stalling, collapse of which also results in a DSB, promotes formation of mtDNA deletions (see for example, Wanrooij et al.^23^ and references therein). In agreement with this observation, the mtDNA sequences in the neighborhood of deletion breakpoints have been shown to be enriched with sequence motifs that might induce mtDNA replication stalling, such as homopolymeric repeats^24,25^, G-quadruplexes^26,27^ and stem-loops (SLs)^28^.

Despite the broad evidence for the involvement of DSBs in initiating mtDNA deletions, the mechanism by which DSBs contribute to mtDNA deletions and, more specifically, which of the DSB repair pathways is involved, has yet to be established. In nuclear DNA, DSBs activate one of several DNA repair pathways, including non-homologous end joining (NHEJ), micro-homology mediated end joining (MMEJ or alternative NHEJ), single strand annealing (SSA), homologous recombination (HR), synthesis dependent strand annealing (SDSA) and break induced replication (BIR)^29^. *Ex vivo* experiments using mitochondrial extracts have demonstrated, to different degrees of confidence, the existence of mitochondrial NHEJ^30–32^, MMEJ^33^ and HR repair pathways^34^. Mitochondrial BIR activity has been previously reported for yeast^35^, but the activity of this repair pathway in mammalian mitochondria has not yet been studied. One of the key features that differentiates these different DSB repair pathways is the length of homologous sequence (HS) around the DSB sites involved in initiating the strand annealing step of the repair process^36–39^. The NHEJ pathway relies on very short HS of 0-5 bp, while the MMEJ and BIR pathways require short (micro) HS of at least 5-25bp^39^. On the other hand, SSA and HR both utilize and require longer HS of ≥30 bp and >100bp, respectively^36,38,40^. In addition, HR requires a template, for example from a sister chromatid, to repair DSBs. The length of the sequence homology around deletion breakpoints has therefore been commonly used as a genomic signature to deduce the repair pathway involved in deletion mutagenesis ^41^. Except for HR, errors in each of the above DSB repair pathways have been shown to cause deletion mutations in nuclear DNA^29^.

In the context of the DSB repair error model, the length of the HSs associated with deletions can provide both discriminating and incriminating information to deduce the specific DSB repair pathway(s) involved in the deletion mutagenesis. In this study, we developed a novel computational method based on DNA hybridization thermodynamics, a mixture distribution model and a maximum likelihood approach, to characterize the involvement of DNA misalignments of various lengths near breakpoints in the formation of mtDNA deletions. More specifically, for a given set of mtDNA deletions, our method gives the composition of misalignment lengths, whose hybridization thermodynamics maximizes the likelihood of observing the observed breakpoint positions. We applied our method to analyze an exhaustive literature compilation of 9,655 deletion breakpoints within the mtDNA major arc region from human, rhesus monkey, mouse and rat. In addition, we sequenced 1,307 novel mtDNA deletion breakpoints of *Caenorhabditis elegans* using next-generation sequencing (NGS). Based on the results of our analysis, we formulated two hypotheses for the formation of mtDNA deletions initiated by DSBs.

## Results

### Mixture Model Formulation

Figures 1 and 2 illustrate the method used in this study to analyse mtDNA deletion breakpoints. The basic assumption of our analytical method is that DNA misalignments play a key role in the formation of mtDNA deletions. Given a set of deletion breakpoints, the method determines the composition of misalignment lengths that maximizes the probability for the nucleotides surrounding the observed breakpoints to participate in a thermally stable DNA misalignment, evaluated using the DNA hybridization thermodynamics. The key steps of the method are briefly described below (for detailed description refer to Online Methods).

**Figure 1.**
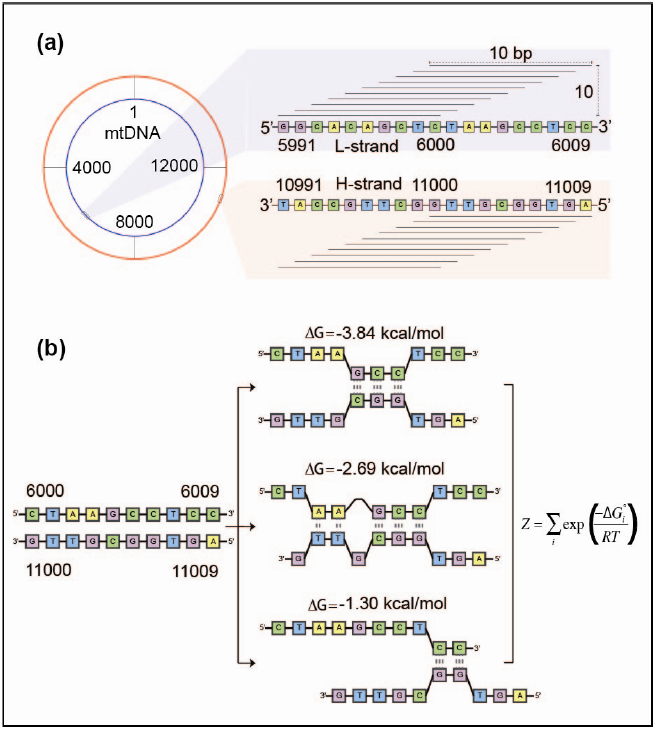
DNA-DNA hybridization partition function calculation. An illustration of the calculation of DNA hybridization partition function for a nucleotide pair at the positions 6000 and 11000 for a misalignment length of 10 bp. (a) A total of 10×10 = 100 distinct pairs of 10 bp sequences can form DNA duplexes that overlap the position pair (6000, 11000) in this illustration. One of the duplex segments comes from the L-strand and the other from the H-strand of mtDNA. (b) For any 10 bp duplex, multiple DNA hybridization configurations could form. For every possible duplex, we computed the hybridization partition function *Z* which reflects the thermal stability of the duplex. The overall hybridization partition function of a given position pair is computed by summing the partition function values of all duplexes overlapping the position.

**Figure 2.**
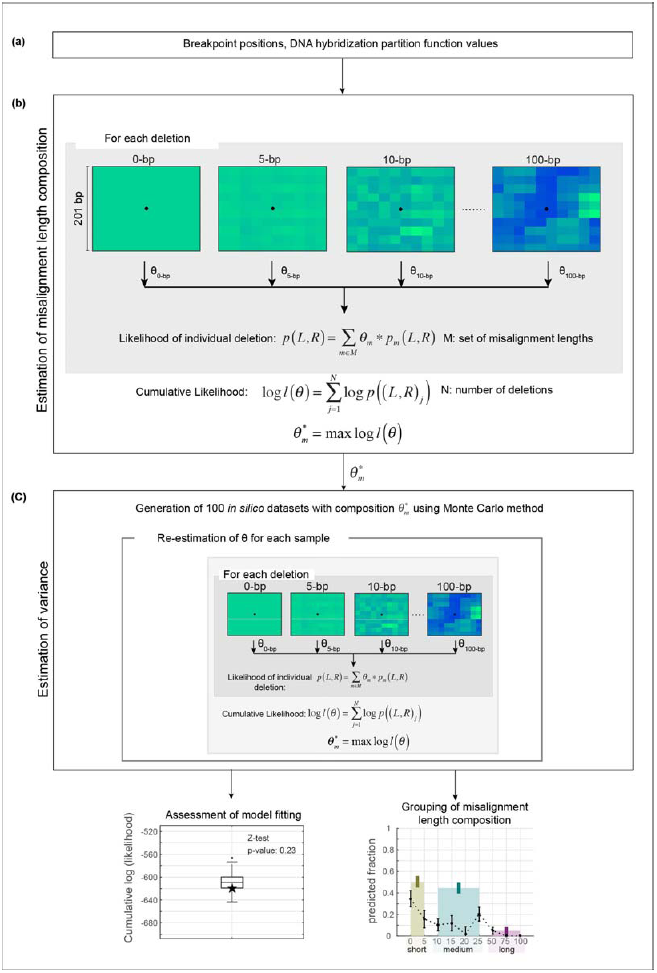
Mixture model analysis of mtDNA deletion breakpoints. (a) We pre-computed the hybridization partition function values (refer Fig.1 and Online Methods) of all feasible nucleotide position pairs between L-strand and H-strand sequence positions containing 5’ and 3’ breakpoints, respectively. (b) For a given mtDNA deletion, we defined a 201 bp × 201 bp window of analysis centered on the breakpoint position (see black asterisk). The analysis window is further divided into bins of 10 bp × 10 bp. The length-specific likelihood (illustrated by the heat maps) indicates the likelihood of sequence pairs in a bin to participate in a DNA misalignment of a particular length, relative to other sequence pairs in the window of analysis. We calculated the length-specific likelihood by dividing the sum of the overall hybridization partition functions of all the nucleotide pairs within a bin with the sum of overall partition functions of all nucleotide pairs in the analysis window. We used the length-specific likelihood of the bin containing the deletion breakpoint as the length-specific likelihood of the mtDNA deletion to arise from DNA misalignment of the corresponding length. The likelihood of the mtDNA deletion was then computed as a linear combination of the length-specific likelihood values from a set of misalignment lengths, according to the mixture distribution model. We determined the unknown misalignment length composition 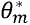 in the linear combination above by maximizing the cumulative likelihood of all mtDNA deletions in a given dataset. (c) For estimating the standard error of 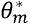, we adopted a Monte Carlo approach by generating 100 *in silico* random breakpoint datasets, running the analysis as described above to each *in silico* dataset, and computing the sample variance of the 100 *in silico θ_m_*’s. For more details, we refer the readers to Online Methods.

For a given mtDNA deletion, we assign a two-dimensional window of analysis consisting of light (L) and heavy (H)- strand nucleotide pairs centered around the breakpoint position (see Fig. 2). We assume that the nucleotide pairs within the window of analysis (one from the L-strand and another from the H-strand) are as accessible for misalignment as the observed breakpoint. For the window of analysis, we extract a 201 bp L-strand sequence centered on the 5’ breakpoint position and a 201 bp H-strand sequence centered on the 3’ breakpoint position (see Fig. 2b). The nucleotide pairs in the window of analysis can have different probability or likelihood to participate in a DNA misalignment, depending the sequence homology and the length of misalignments considered. We use DNA hybridization thermodynamics to compute the length-specific likelihood value for each nucleotide pair in the window of analysis, relative to the neighbouring pairs, to be involved in a DNA misalignment of a given length (see Fig. 1 and Online Methods). A misalignment with a lower free energy of hybridization is more thermally stable and thus has a higher likelihood to form according to the Maxwell-Boltzmann statistics^42^. Here, the DNA misalignment length involved in the formation of a mtDNA deletion is the unknown variable of interest. The length of the most likely contributing sequence is expected to be different from one mtDNA deletion to another, and thus one expects a mixture or composition of misalignment lengths for any set of mtDNA deletions.

In our analysis, the length of DNA sequences hypothesized to take part in misalignments near the breakpoints spans from short DR motifs to 100 bp long duplexes^15,43^. We include misalignment lengths between 0bp and 100bp, specifically 0, 5, 10, 15, 20, 25, 50, 75 and 100bp. The 0bp length accounts for a misalignment-independent (sequence independent) formation of mtDNA deletions. For the calculation of 0-bp length likelihood, we utilize the uniform distribution, instead of the DNA hybridization partition functions, following the principle of equal *a priori* probability^44^. For each misalignment length, we further divide the window of analysis into 10bp × 10bp bins, and assigned the length-specific likelihood of each deletion to that of the bin containing the breakpoint (see Online Methods). The length-specific likelihood value of a bin is set to the sum of the length-specific likelihoods of all nucleotide pairs in the bin. The binning strategy is implemented since the deletion breakpoint may occur in the proximity of – not within – the DNA misalignment.

We utilize a mixture model to evaluate the misalignment likelihood of a deletion as a linear combination of the length-specific likelihood values corresponding to the aforementioned set of lengths (see Fig. 2). Finally, we determined the optimal composition 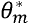 of the misalignment lengths (i.e. coefficients for each misalignment length used in the model) by maximizing the cumulative likelihood of observing a given set of mtDNA deletions (i.e. by summing the likelihood values of all deletions in the dataset; see Fig. 2). In this study, we interpreted the optimal composition 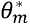 of a specific length as the fraction of mtDNA deletions in the given set that form by way of DNA misalignments of that length. In interpreting the result of the analysis, we grouped the optimal composition 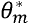 into three length categories: short 0-5 bp, medium 10-25 bp, and long 50-100 bp.

To test if our method is able to recover the DNA misalignment length composition in cases where the true composition is known, we generated *in silico* nucleotide position pairs as benchmark datasets using Monte Carlo sampling for several pre-specified length compositions (see Online Methods). The analysis of the benchmark *in silico* datasets demonstrated that the ability of our method to accurately recover the pre-specified misalignment length composition used in the *in silico* data generation (see Supplementary Methods and Supplementary Fig. 1). In addition, for each of the real dataset used this study, we performed our entire analysis using different window and bin sizes to test for any dependence of the results on these parameters. We found a high degree of correlation among runs using widely varying parameters. Thus, the general findings of our study did not sensitively depend on the sizes of the window of analysis and the binning (see Supplementary Fig. 2).

### Mammalian mtDNA deletions predominantly exhibit short and medium length misalignment signatures near breakpoints

For our main analysis, we compiled an exhaustive compendium of 9,655 mtDNA deletion breakpoint positions previously reported in human, rhesus monkey, mouse and rat, and categorized these deletions into 15 groups based on the species and the experimental conditions (patient groups used, mutant strains, treatment conditions; for more details see Table 1). This compendium comprised, to the best of our knowledge, all mutant sequences publicly available at the time of this study. The first class of mtDNA deletions that we analyzed were those reported in transgenic mice expressing mitochondria-targeted restriction enzymes PstI and ScaI (see Fig. 3a)^19–21^. Overexpression of restriction endonucleases introduces DSBs at specific restriction sites in the mtDNA sequence and results in mtDNA deletions typically with one or both breakpoints located in direct proximity to these restriction sites. Since this class of deletions derived from DSBs, the mtDNA deletion breakpoints represented a good case to investigate the misalignment length distribution associated with DSB-induced mtDNA deletions. The outcome of our analysis shows a significant fraction of short misalignments (0 – 5 bp), a minor contribution from medium length (10 – 25 bp), and practically no participation by longer misalignments (≥ 50 bp). The significance of short misalignments is indicative of a key role of non-homologous and micro-homology end-joining (NHEJ/MMEJ) repair mechanisms in the creation of mtDNA deletions caused by restriction enzyme-induced DSBs. On the other hand, the absence of contribution from long misalignments signifies a lack of involvement from homologous recombination repair pathway in the mutagenesis of these deletions.

**Figure 3.**
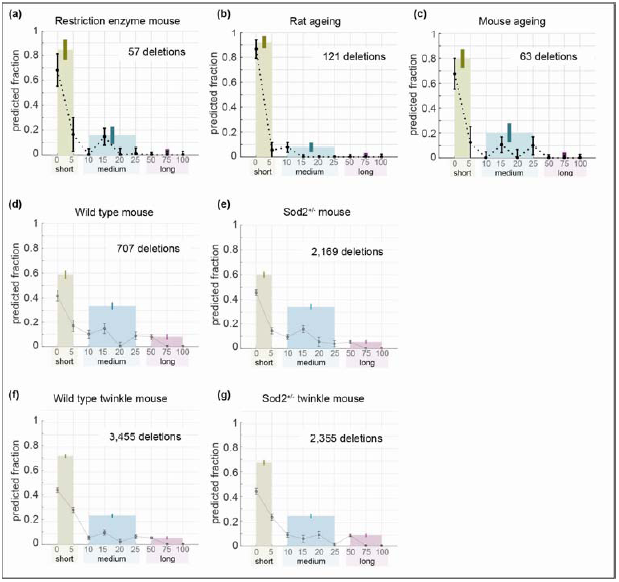
Analysis of mtDNA deletion breakpoints from rodents. Mixture model coefficients for (a) transgenic mice with mitochondrial restriction enzymes, (b) rat ageing, (c) mouse ageing, (d) wild type mice, (e) Sod2^+/−^ mice, (f) wild type mice with Twinkle overexpression and (g) Sod2^+/−^ mice with Twinkle overexpression. The colored bars show the sum of the coefficients for short (0 – 5 bp), medium (10 – 25 bp) and long (≥ 50 bp) duplexes. The error bars indicate the sample standard deviation estimated using 100 *in silico* datasets (see Online Methods).

**Table 1:**

| Dataset count | Species | Dataset Name | Total NR deletions | Source/Description |
| --- | --- | --- | --- | --- |
| 1 | Human | Mesial temporal lobe epilepsy | 208 | Epilepsy/sclerosis patient brain samples with chronic inflammation |
| 2 |  | Ageing | 151 | Aged tissues: muscle, brain, skin |
| 3 |  | Myopathy | 138 | KSS/CPEO, non-autosomal, primarily skeletal muscle |
| 4 |  | Autosomal Myopathy | 85 | CPEO, autosomal, mutation in POLG gene |
| 5 |  | Myositis | 49 | Inflammation in muscle |
| 6 |  | Pearson Syndrome | 34 | Rare childhood disorder, primarily blood cells |
| 7 |  | Charcot Marie Tooth Disease | 31 | Inherited nervous disorder, can onset early |
| 8 | Rat | Ageing | 121 | Aged tissues |
| 9 | Mouse | Ageing | 63 | Aged tissues, only within limited region of major arc |
| 10 |  | DSB | 57 | Deletions induced by Pst1 and ScaI enzymes |
| 11 |  | WT | 707 | Wild type control mice |
| 12 |  | Sod | 2,169 | Superoxide dismutase (Sod2<sup>+/-</sup>) heterozygous mutant mice |
| 13 |  | WT;TW+ | 3,455 | Wild type mice with TWINKLE overexpression |
| 14 |  | Sod;TW+ | 2,355 | Sod2<sup>+/-</sup> heterozygous mice with TWINKLE overexpression |
| 15 | Rhesus Monkey | Ageing | 32 | Aged tissues |
|  |  | Total | 9,655 |  |

We subsequently analyzed three groups of mtDNA deletions found in wild-type (WT) rats and mice without any transgenic expression or diseases (Figs. 3b – d). The rat dataset and one of the mouse datasets comprised mtDNA deletion breakpoints gathered from different reports in the literature (see Supplementary Document 2). The second and larger mouse dataset included mtDNA deletion breakpoints from a single NGS study^45^. The results of our analysis of these datasets, as depicted in Figures 3b – d, show misalignment length compositions that resembled that of DSB-induced mtDNA deletions above. Again, the largest fraction of the deletions is associated with short misalignments, with the medium lengths having a lower contribution and the long misalignments having the lowest fraction. The similarity in the misalignment characteristics to DSB-induced mtDNA deletions supports an important role of DSBs in the formation of mtDNA deletions in WT mice and rats.

The aforementioned NGS study also generated mtDNA deletion breakpoints for mice heterozygous for *sod2* gene expression. Mitochondrial superoxide dismutase (Sod2) converts superoxide anions (O_2_^−^) generated from electron transport chain into hydrogen peroxide, and a complete lack of Sod2 (homozygous Sod2^−/−^) results in early postnatal lethality in mice^46,47^. Meanwhile, heterozygous Sod2^+/–^ mice suffered increased mitochondrial oxidative damage, higher burden of mtDNA point mutations, more frequent mtDNA replication stalling and elevated levels of recombined mtDNA molecules^48^. The breakpoint hotspots also coincided with G-rich regions in the mtDNA H-strand, suggesting increased oxidative damage (8-oxoG) as the causative factor of higher levels of mtDNA deletions in the Sod2^+/–^ mice^45^. Our analysis of mtDNA deletion breakpoints in these Sod2^+/–^ mice (Fig. 3e) suggests a misalignment length distribution that is similar to that of WT mice (Figs. 3c and d). Consistent with the above finding, our comparison of the breakpoint position distributions shows that the mtDNA deletion hotspot regions in Sod2^+/–^ mice coincides with those in WT mice (see Supplementary Fig. 3). Taken together, while a reduced capacity of Sod2 increases the frequency of mtDNA deletions, the similarity in the misalignment length compositions and the deletion hotspots between WT and Sod2^+/–^ mice suggests that oxidative damage contributes significantly to the mtDNA deletions in WT mice. The likeness in the misalignment length compositions of WT and Sod2^+/–^ mice to those of PstI and ScaI mice further supports the involvement of DSBs in mtDNA deletion formation induced by oxidative damage.

The same NGS study further showed that the overexpression of TWINKLE helicase (TW^+^) in Sod2^+/–^ mice rescued the oxidative stress phenotype^48^. While the frequency of mtDNA rearrangements, including deletions, in Sod2^+/–^;TW^+^ mice was lower than in Sod2^+/–^ mice, it remained elevated in comparison to WT. Indeed, TWINKLE overexpression in WT mice led to an increased level of mtDNA rearrangements (see Supplementary Figure S6 in the original publication^48^). In addition to its putative function as a helicase, TWINKLE has been shown to catalyze DNA recombination^49,50^. A recent study further demonstrated that strand exchange activity of TWINKLE requires 3-6 bp sequence homology^50^. Thus, the increased frequency of mtDNA rearrangements in WT;TW^+^ mice may arise from higher DNA recombination activity due to TWINKLE overexpression. The misalignment length compositions of mtDNA deletions in WT;TW^+^ and Sod2^+/–^;TW^+^ mice mirror each other (see Figs. 3f and g). But, in comparison to WT and Sod2^+/–^ mice, mtDNA deletions of WT;TW^+^ and Sod2^+/–^;TW^+^ mice have a higher fraction of the short length misalignments, specifically 5bp. Thus, our analysis is sensitive enough to detect the higher frequency of short 5bp misalignment that is expected from the overexpression of TWINKLE in the above mice.

Our analysis of mtDNA deletions found in aged rhesus monkey and human shows misalignment length compositions similar to that of rodent mtDNA deletions, albeit with nominally higher fractions of medium length misalignments in the primate datasets (see Figs. 4a and b). In addition to ageing, mtDNA deletions in human have also been commonly associated with sporadic and autosomal myopathies and several other neuromuscular and multi system disorders. We compiled human mtDNA deletions reported for patients with neuromuscular disorders from the literature, and categorized them into 6 disease groups based on the patient description in the source articles (see Online Methods). Except for mtDNA deletions associated with polymerase gamma (*polg*) gene mutations, our analysis of these groups of mtDNA deletions gives compositions of misalignment lengths that are similar to that of age-related mtDNA deletions in human (see Figs. 4c – g). Thus, human mtDNA deletions during ageing and in neuromuscular diseases (except autosomal *polg* mutations dataset) appear to occur by similar mechanisms.

**Figure 4.**
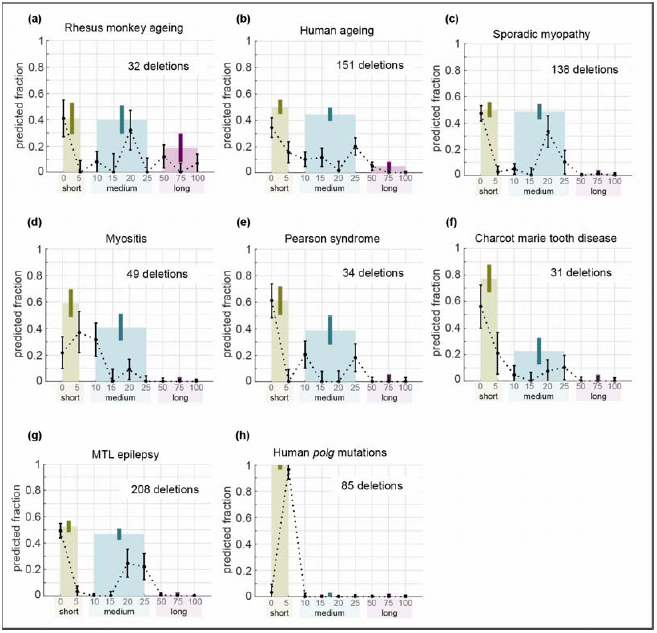
Analysis of mtDNA deletion breakpoints from primates. Mixture model coefficients for (a) rhesus monkey ageing, (b) human ageing, (c) sporadic myopathy, (d) myositis, (e) Pearson’s syndrome, (f) Charcot Marie tooth disease, (g) mesial temporal lope (MTL) epilepsy and (h) myopathy patient with compound *polg* gene mutations. The colored bars show the sum of the coefficients for short (0 – 5 bp), medium (10 – 25 bp) and long (≥ 50 bp) duplexes. The error bars indicate the sample standard deviation estimated using 100 *in silico* datasets (see Online Methods).

The one exception to this pattern is the autosomal *polg* mutations. POLG is the sole DNA polymerase present in mammalian mitochondria. Mutations in *polg* gene cause defects in the mtDNA replication process, leading to mtDNA depletion and mutations, including deletion and point mutations^25,51^. The associated mtDNA deletions have breakpoints near HPs and GQs, sequence motifs that are known to induce replication stalling^25,26^. Our analysis of mtDNA deletion breakpoints from autosomal myopathy patient with compound *polg* mutations, indicates that these mtDNA deletions are associated with only short HSs (see Fig. 4h). The lack of involvement of medium and long HS in mutant POLG dataset is in agreement with a previous study demonstrating a role of POLG in misalignment-dependent DSB repair in human^52^. More specifically, the study showed that the common 4,977 bp mtDNA deletion in human could be induced artificially by DSBs, through a mechanism that requires a functional POLG. In addition, other DNA polymerases, such as Pol-θ, have previously been shown to directly take part in micro-homology (i.e. medium length) mediated end joining DSB repair in nuclear DNA^36^.

In summary, our analyses of mtDNA deletion breakpoints from four mammalian species, including human, point to a shared mtDNA deletion mutagenesis mechanism, involving predominantly DNA misalignments of short and medium HS lengths (0-25bp). In the context of DSB repair, the significance of short to medium HS misalignments is indicative of erroneous NHEJ and/or MMEJ pathways. Meanwhile, DNA misalignments of >25bp appear to have little or no involvement in the formation of mtDNA deletions. Except for PstI and ScaI transgenic mice dataset, the damage that initiates the formation of mtDNA deletions for other datasets used in our study is unknown and likely arises from multiple sources. However, the similarity of the misalignment length compositions for PstI and ScaI mice, WT mice and rats, and Sod2^+/−^ mice, gives support to the involvement of DSBs and oxidative damage in the deletion mutagenesis in rodents.

### Ageing associated mtDNA deletions in nematode *C. elegans*

As the analysis of mammalian mtDNA deletions above provided evidence for a shared mutagenesis mechanism among the mammals, we carried out an analysis using mtDNA deletions from a different phylum, more specifically *C. elegans.* The short life span of *C. elegans* and their amenability to genome scale genetic manipulations make the nematode a powerful model organism to study the biology of ageing, including the mechanisms of mtDNA deletion formation. Similar to mtDNA deletions reported in mammals^13^, a small number of mtDNA deletions and breakpoints have been reported in *C. elegans* and these tend to be flanked by DRs^53,54^. More specifically, 5 out of 6 known age-related mtDNA deletions sequenced in *C. elegans* have DRs flanking their breakpoints^53,54^. In this study, we used NGS to generate a much larger dataset, identifying 1,307 unique age-related mtDNA deletion breakpoints in *C. elegans* sampled at three different ages: days 4 (371 deletions), 7 (416 deletions) and 10 (520 deletions). Consistent with previous reports^53,54^, a large fraction (82%) of these mtDNA deletions were flanked by DRs of ≥3 bp. Our analysis of the mtDNA deletion breakpoints in *C. elegans* suggests that short and medium misalignments are most important with only a negligible involvement of lengthy misalignments (see Fig. 5). This observation is consistent with our findings from rodent and primate mtDNA deletions. The results from nematodes from different age groups differ only slightly. Thus, age-related mtDNA deletions in *C. elegans*, like those seen in mammalian species (see above), are associated mainly with short and medium HS misalignments, indicating that the mechanism of mtDNA deletion mutagenesis appears to be conserved across distant taxonomic groups.

**Figure 5.**
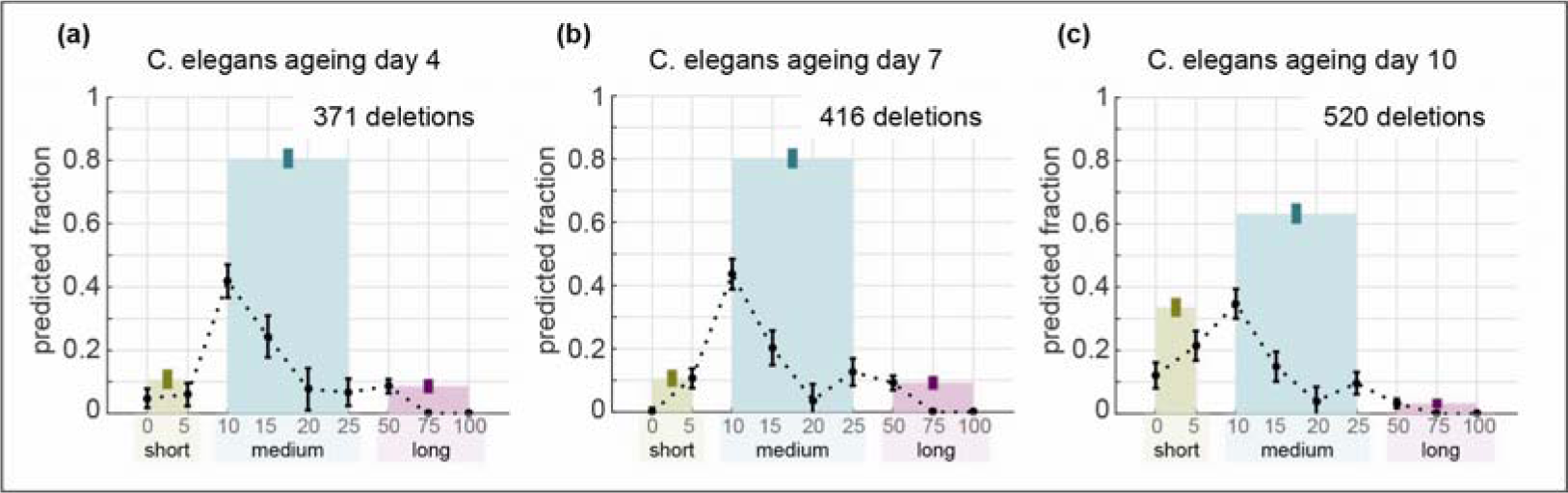
Mixture model predictions for mtDNA deletions in nematode *C. elegans*. Mixture model coefficients predicted for age-related mtDNA deletion datasets from *C. elegans* at (a) Day 4, (b) Day 7 and (c) Day 10. The colored bars show the sum of the coefficients for short (0 – 5 bp), medium (10 – 25 bp) and long (≥ 50 bp) duplexes. The error bars indicate the sample standard deviation estimated using 100 *in silico* datasets (see Online Methods).

## Discussion

The formation and clonal expansion of mtDNA deletions cause mitochondrial respiratory dysfunction, which in human leads to a class of clinically heterogeneous disorders collectively known as mitochondrial diseases. Over the past decades, experimental and mathematical modeling studies have provided a better understanding of the etiology of mtDNA mutation expansion^55–59^. In contrast, the origin and mechanism of mtDNA deletions are less well understood. In the literature, there exist broad empirical evidence from human and model organisms that DSBs are involved in the etiology of mtDNA deletions and that deletions may result from erroneous DSB repair^12^. However, the specific DSB repair pathway(s) involved and the mechanism by which the corresponding errors cause mtDNA deletions have yet to be established. Here, we developed a novel method that combines DNA hybridization thermodynamics, a mixture distribution model and maximum likelihood estimation, for determining the lengths of DNA misalignments associated with mtDNA deletion breakpoints. We used the misalignment lengths as a signature for identifying the specific DSB repair errors involved in mtDNA deletion formation and to examine similarities and differences in the mutagenesis mechanisms of mtDNA deletions from different species and conditions.

Our analysis of mtDNA deletions across five species (human, rhesus monkey, rat, mouse and nematode worms) suggests that the deletion breakpoint positions are most consistent with a mutagenesis mechanism that is driven by the thermodynamics of short to medium length DNA misalignments (0-25 bp). The contributions from longer misalignments vary slightly among species and conditions, but in general, are low. Assuming mtDNA deletions indeed arise by errors during DSB repairs, then the significant combined contribution of short and medium DNA misalignments implicates NHEJ and micro homology dependent mechanisms (e.g. MMEJ). NHEJ is generally known as the quickest and the most common route of repairing DSBs *in vivo*^60^. In comparison to NHEJ, MMEJ repair has slower kinetics, increasing the chance of chromosomal rearrangements caused by unrepaired DSBs^60^.

At the same time, the low contribution from long misalignments excludes HR as a main source of mtDNA deletions. The lack of evidence for the involvement of HR in mtDNA deletion formation is in good agreement with two well-known aspects of the HR repair in mitochondria. First, HR is generally considered an error-free mechanism as the repair uses a template DNA molecule^29^. Second, individual mtDNA molecules are typically isolated into separate nucleoids, limiting inter-molecular recombination that is necessary for HR repair^61^. While our analyses do not exonerate replication slippage as a mechanism of deletion formation, one expects higher contributions from longer and more thermodynamically stable misalignments, if deletions are to occur predominantly by the slip-strand mechanism^62^.

In contrast to our finding on the lack of involvement of lengthy misalignments, a previous study by Guo *et al.* using mtDNA deletion breakpoints from human frontal cortex found that the breakpoints are over-represented in regions of mtDNA sequences that can form highly stable 100-bp misalignments^43^. However, while Guo *et al.* and our study were both based on the same DNA hybridization thermodynamics, our analysis differed from that in Guo *et al.* in two major aspects; (1) we used the thermodynamic partition function values to account for the thermal stability of all feasible duplex configurations based on the Maxwell-Boltzmann statistics^42^, whereas Guo *et al*. relied on only the duplex configuration with the minimum free energy value; (2) we took into consideration the DNA misalignments of different lengths, whereas Guo *et al*. considered only 100bp long misalignments. When we repeated the analysis of Guo *et al*. using the same breakpoints but allowing for shorter misalignment lengths, more specifically 5, 10, 25, 50 and 100 bp, we observed over-representation of deletion breakpoints in regions of mtDNA forming stable duplexes for each of these lengths, suggesting that there is, in fact, no contradiction between the two studies insofar as both methods identify the same sequences (see Supplementary Fig. 4 and Supplementary Methods).

There still exists an obvious inconsistency between the 2 – 25 bp short deletions commonly caused by NHEJ/MMEJ errors, and the 1000s of bp long deletions observed in mtDNA. Despite the rarity of large deletions resulting (>1000bp) from erroneous NHEJ/MMEJ repair, studies of nuclear DNA have shown a scenario involving multiple DSBs by which these repair errors could cause such deletions^63,64^. We therefore posit that multiple DSBs (at least two) on a single mtDNA molecule could initiate the formation of a large deletion, where the DNA ends from two distinct DSBs are misjoined by NHEJ/MMEJ repair (see Fig. 6a). Here, any two distantly located DSBs on a mtDNA molecule might come into a close proximity to each other because of the folding and packing of mtDNA by mitochondrial transcription factor A (TFAM)^65^. Alternatively, the misjoining of distant DSB ends can occur if the mtDNA sequence in between these ends is degraded. The recombination between two mtDNA molecules, each with a DSB, may also result in a deletion mutation^19^. But, such events have been shown to take place only very rarely *in vivo*^19^ and is therefore unlikely to contribute significantly to mtDNA deletion formation under physiological conditions.

**Figure 6.**
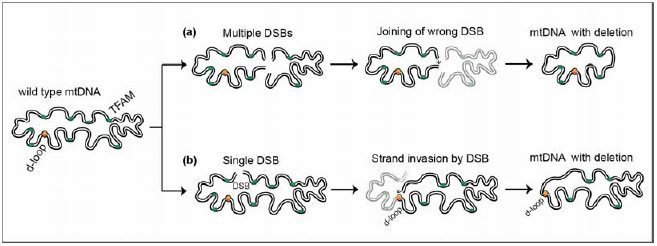
DSB-induced mtDNA deletions. Mitochondrial DNA are folded and packed into nucleoids with the help of TFAM protein molecules. Due to the compact organization of mtDNA, physically distant positions in a mtDNA molecule could come into close proximity. (a) When multiple DSBs occur at such positions, the misjoining of DSB ends from distinct DSBs could create a deletion mutation. (b) Alternatively, one end of a DSB could invade and recombine with an open region in the mtDNA, such as the D-loop, producing a deletion mutation.

Is there an alternative explanation where a single DSB could cause large mtDNA deletion? In order to explain the D-loop hotspot among mtDNA deletions found in the PstI transgenic mice, Srivastava et al.^21^ proposed that one of the ends of the DSB at the restriction site recombines with the open D-loop structure. Extending this proposed mechanism, we hypothesize that a DSB end could invade any open structures in mtDNA, such as the D-loop, RNA-loop (R-loop)^66^ or TFAM bubbles^67^ (see Fig. 6b). The natural question that follows from this hypothesis is whether there exists a possible mitochondrial protein that is able to assist the strand invasion. Recent studies have demonstrated that TWINKLE helicase possesses several DNA modifying capacities including strand invasion (or strand-exchange)^49,50^. Consistent with the importance of short to medium HSs in our analyses, the efficiency of the strand-exchange assisted by TWINKLE depends strongly on the sequence complementarity of the first 3-6 bp of the invading strand^49,50^.

The two mechanistic hypotheses of DSB-induced mtDNA deletions are consistent with the presence of mtDNA deletions in PstI and ScaI transgenic mice with breakpoints located near the restriction enzyme cleavage (DSB) sites and with the well-known breakpoint hotspot in the D-loop that has an open structure, making it vulnerable to both DSBs and strand invasion. The D-loop, region around mtDNA deletion breakpoints have also been reported to be enriched with sequence motifs, such as GQs^26^, HPs^25^ and stem loops^28^, that are capable of inducing DSBs by stalling and collapse of replication fork. However, our hypotheses do not directly explain the presence of DRs in the vicinity of the deletion breakpoints, as commonly observed.

By generating *in silico* deletion breakpoint datasets using the misalignment length compositions from the analyses of WT human and mouse datasets, we found that the *in silico* mtDNA deletions have breakpoints that are as near to a DR as the reported mtDNA deletions (see Supplementary Fig. 5). Thus, the deletion mutagenesis mechanisms, involving short and medium length misalignments, proposed in this study are able to explain the commonly observed features of mtDNA deletions.

## Methods

### Deletion breakpoint positions and mitochondrial DNA sequences

We compiled 9,655 breakpoint positions of mtDNA deletions from published literature and grouped them into species- and clinical condition-specific categories (see Supplementary Document 2). Deletions reported in normal aged tissues from human, rhesus monkey, mouse and rat were grouped under age-related deletions. Disease-associated human mtDNA deletions were further grouped under sporadic myopathy (including Kearns-Sayre syndrome (KSS) and chronic progressive external ophthalmoplegia (CPEO)), myopathy with compound mutations in *polg* gene, Pearson syndrome (PS), Myositis, Mesial temporal lobe epilepsy and Charcot Marie tooth disease based on the clinical description of the patients. Our datasets also included mtDNA deletions from mice expressing mitochondrial Pst1/Sca1 restriction enzymes, Sod2^+/−^ heterozygous mice, wild type control mice and Twinkle overexpression mice. In this study, we have considered only the deletions occurring within the mtDNA major arc region in mammals, as deletions involving the minor arc were much less frequent. Within each group of mtDNA deletions, identical breakpoint positions were accounted only once.

In the analyses, we used the complete, annotated mtDNA sequences for human (NC_012920), rhesus monkey (NC_005943), mouse (NC_005089), rat (X14848) and nematode worms (NC_001328) from NCBI database. The breakpoint positions were reported based on the L-strand nucleotide sequence.

### C. elegans strains and maintenance

The strain JK1107 (*glp-1*) was cultivated and maintained at 25 °C to prevent progeny. Preparation of nematode growth media was done as previously described^68^ with the addition of Streptomycin (Sigma) at the final concentration of 200 μg/ml. Streptomycin-resistant bacteria strain (*Escherichia coli* OP50-1) was added to the solid media at 10^10^ cells/ml.

### Extraction and purification of mtDNA

Adult worms were harvested on day 4 (young), 7 (middle-aged) and 10 (old). Worms were washed off plates using ice-cold M9 buffer^68^ and allowed to settle on ice, washed twice with ice-cold S-basal buffer to remove bacteria, and twice with ice-cold isolation buffer (210 mM mannitol, 70 mM sucrose, 0.1 mM EDTA, 5 mM Tris-HCl, pH 7.4). Worms were homogenised in isolation buffer on ice, using a Teflon pestle homogeniser. Homogenate was sampled and checked under light microscope to ensure complete homogenisation. Worm homogenate was centrifuged at 600 g, 4 °C for 10 min to remove cell debris. Supernatant was centrifuged at 7,200 g, 4 °C for 10 min to obtain mitochondria. Pellet was resuspended in TE buffer (50 mM Tris-HCl, 0.1 mM EDTA, pH 7.4). mtDNA was extracted and purified using PrepMan Ultra Sample Preparation Reagent (Thermofisher Scientific) according to the manufacturer’s protocol.

### NGS Sample Preparation

An approximately 6.3kB PCR product (positions 1800 – 8100 in *C. elegans* mtDNA) from purified mtDNA was fragmented to an average size of 350 base pairs using Covaris sonication (Model S2), starting with 100-200ng of DNA. Paired end libraries were prepared using the TruSeq Nano DNA Library Prep Kit (Illumina) or NEBNext Ultra DNA Library Prep kit for Illumina (New England Biolabs, U.S.A). Each library contained a unique barcode for demultiplexing following sequencing. Libraries were analyzed for size distribution and quality using a DNA high sensitivity chip on a Bioanalyzer 2100 instrument (Agilent). Libraries were quantified by qPCR (Kapa Biosystems), normalized and equally pooled for sequencing. Sequencing was performed on an Illumina miSeq using a 2x150bp sequencing kit. Fastq files were generated on-board the miSeq. The quality of the data was confirmed using FastQC prior to bioinformatic analysis.

### Analysis of probability of DNA-DNA misalignments within mtDNA major arc

In the analysis of mtDNA deletion breakpoints, we used DNA hybridization thermodynamics to assign the length-specific likelihood of a particular nucleotide pair, one from the L-strand and another from the H-strand, to be involved in a misalignment of a particular sequence length. More specifically, for a given pair of DNA positions (*l,r*), where *l* is a nucleotide position in the L-strand and *r* is a nucleotide position in the H-strand, and for a chosen DNA sequence length *m*-bp, we first identified all *m×m* DNA duplexes overlapping the position (*l,r*) (Fig. 1a). We then computed the partition function of DNA-DNA hybridization (*Z*) for every duplex above. This partition function is defined by

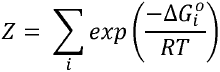

where, 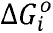 is the Gibb’s free energy of hybridization of the *i^th^* conformation^69^. Note that multiple conformations could exist between a given pair of hybridizing duplexes (Fig. 1b). The partition function thus sums up the probabilities of occurrence of all possible duplex conformations that could form between the given pair of DNA segments. Here, we employed the *hybrid* subroutine of the UNAFold package, setting the temperature to 37° C for mammalian datasets and 25.5° C for *C. elegans* datasets, while keeping the default values for all other parameters^70^. Provided the partition function values for every duplex, we computed the overall partition function of the position (*l,r*) denoted by Z*m*(*l,r*) as follows:

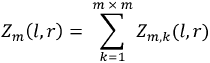

where *Z*_*m,k*_(*l,r*) is the partition function of the *k*^th^ DNA sequence pair associated with the position *(l, r)*.

To compute the length-specific likelihood, we defined a window of the analysis. The window corresponded to 201bp × 201bp region centered on each breakpoint position (i.e. ± 100 bp sequence from the breakpoint (*l, r*) position). We defined the window by (*L*_*w*_, *R*_*w*_), where *L*_*w*_ and *R*_*w*_ denote the sets of the left and right mtDNA nucleotide positions in this window. We further binned the overall partition function values of neighboring mtDNA positions within the window of analysis. Unless specifically mentioned, a standard bin dimension of 10 bp × 10 bp was used throughout this study. We denoted each bin by (*L, R*), where *L* and *R* describe the ranges of the left and right mtDNA positions inside the bin, respectively. Finally, for a given deletion breakpoint, we evaluated the length-specific likelihood or probability of DNA misalignment *p_m_*(*L, R*) for the bin (*L, R*) containing the breakpoint position, according to

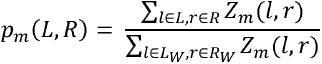

### Mixture model analysis

We formulated a mixture distribution model to estimate the composition of misalignment lengths in a given set of mtDNA deletions. The estimation was based on maximizing the cumulative likelihood of the deletions in the dataset. In the mixture model, the likelihood of a deletion was given by a linear combination of the length-specific likelihood values of the deletion from different sequence lengths. In particular, for a given bin (*L, R*) containing a deletion breakpoint position pair, we calculated the following mixture probability

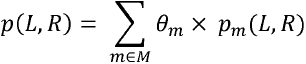

where M denotes the predefined set of sequence lengths and *θ*_*m*_ denotes the fractional contribution of the sequence length *m*-bp. We used the following set of sequence lengths *M* = {0, 5,10,15,20,25,50,75,100} in this study. Finally, we carried out a maximum likelihood estimation to find the optimal composition 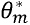 that maximizes the cumulative likelihood, as follows:

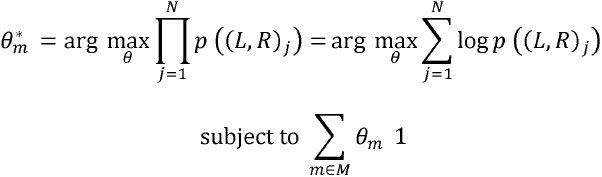

where (*L, R*)_*j*_ corresponds to the *j*^th^ mtDNA deletion breakpoint and *N* denotes the number of mtDNA deletions in the analysis. The optimization was performed using *fmincon* function in MATLAB (version 2015a; MathWorks, Inc.) with a multi-start strategy using 5 different starting points to ensure that we obtained a globally optimal solution.

In order to provide the variance of the optimal 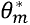, we generated 100 *in silico* breakpoint datasets using the estimated 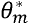 (see below for details) and repeated the mixture model analysis on each of the *in silico* dataset. The variance of 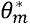 was set equal to the sample variance from the results of the analyses of 100 *in silico* datasets. We also compared the optimal (log-) likelihood value of the mtDNA deletion breakpoint analysis with the distribution of (log-) likelihood values from the *in silico* datasets. In general, the optimal likelihood values from the analysis of mtDNA deletion datasets were not significantly different from the optimal likelihood values from the *in silico* datasets generated using the same optimal 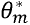, indicating that the mixture distribution model was able to describe the likelihood of the observed mtDNA breakpoint positions as well as that of the *in silico* generated breakpoints.

### Generation of *in silico* breakpoints data

*In silico* breakpoint set is a set of breakpoint pair positions that are computationally generated by random sampling from the likelihood values over given window(s) of analysis. The *in silico* dataset exhibit a specific misalignment signature at their breakpoint positions by setting the composition 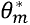. To generate an *in silico* breakpoints dataset we need (1) 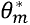, (2) the length-specific likelihood values for mtDNA sequence basepair positions, (3) the sample size, (4) the window(s) of analysis. From 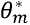and the sample size, we first calculated the number of breakpoint pairs to be sampled from each HS length component. Within a given window of analysis, we evaluated the length specific likelihood of DNA misalignment *p*_*m*_(*L, R*) by following the procedure described above. *In silico* breakpoint pair positions are randomly sampled from within the window of analysis using an inverse sampling procedure with *p*_*m*_(*L, R*) as the propensity^71^. In the calculation of variance of 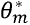from the analysis of reported mtDNA deletions, the analysis windows of the original dataset were used in generating *in silico* deletion breakpoints. Here, for each random *in silico* breakpoint, we chose randomly an analysis window from those of the original dataset without replacement.

### Statistical analysis

Statistical analyses were performed using in built functions in MATLAB (version 2015a; MathWorks, Inc.). Two-sided z-test was used for statistical comparison of log -ikelihood values between dataset and *in silico* generated samples. Two-sided *t*-test and Mann Whitney U (MWU) test were used for statistical comparison of mean, median values of breakpoint DR distance values, respectively. Unless stated otherwise, a statistical significance is set at p-value < 0.05.

## Acknowledgements

We thank Dr. Jaakko Pohjoismäki and Dr. Sion Williams for kindly providing their mouse mtDNA deletion breakpoints dataset for our analysis.

